# Revealing Spatial Heterogeneity across the Gut Tissue-Lumen Interface through MALDI Mass Spectrometry Imaging

**DOI:** 10.1101/2025.06.01.657309

**Authors:** Jacob J. Haffner, Soo Hyun Ahn, Tian (Autumn) Qiu

## Abstract

Metabolites play critical roles in modulating gut-microbe interactions and are closely related to health and disease consequences. One critical aspect of the gut-microbe interactions is spatial heterogeneity, particularly at the interface space between gut tissue and luminal contents, where a unique microhabitat of diverse microbial functions encounters the host tissues. Exploring the spatial heterogeneity of this interface enables insights into these gut-microbe interactions. Previous studies commonly investigated metabolome through tissue homogenization and bulk analysis, but these methods result in the loss of spatial information. In this project, we use high-resolution matrix-assisted laser desorption/ionization mass spectrometry imaging (MALDI-MSI) approaches to reveal the chemical spatial heterogeneity of colon tissue, luminal contents, and the tissue-lumen interface across multiple colonic regions. We applied a swiping technique to aid the preservation of luminal content integrity and used two MALDI matrices to cover a wide range of metabolites. The MALDI-MSI analyses revealed distinct patterns of metabolite spatial localization across the gut tissue-lumen interfaces, amongst which the interface-enriched features are of particular interest due to their possibly connection to the gut-microbe interactions. Overall, the rich spatial heterogeneity of metabolomic profiles across the gut tissue-lumen interfaces highlight the molecular participants in host-microbe interactions, providing new opportunities for examining host-microbiome-metabolome dynamics.

## Introduction

The mammalian gastrointestinal tract (GI tract) houses complex ecosystems of microbiota whose functions and distribution heavily inform host health^1,2^. Such microbially-derived effects for the host include immunity boosts^3,4^, energy metabolism^5–7^, pathogen resistance^8,9^, food digestion^10,11^, influences on health and disease states^12–15^, to name a few. The GI tract represents one of the largest microbially-interfacing sites in the human body for interactions between the host, their resident microbes, and the environment^1^. Small molecules, called metabolites (<1500 Da^16–18)^, comprises the network of chemical interactions known as the metabolome, and play a central role of host-microbe interactions^17,18^. Examples of key metabolites in such interactions include bile acids and short chain fatty acids, both associated with host health and disease states^14,19–23^. Specific dysbiosis states, like gut inflammation and colitis^21,24,25^, and their host interactions in specific areas of the gut such as the colon are linked to specific subsets of microbial metabolites An in-depth characterization of host-microbe metabolism can facilitate the identification of key metabolites and pathways that may serve as targets for microbiome-based disease treatment.

The spatial localization of metabolites is highly relevant for their biological functions ^26–28^. Host tissues and microbiome form interkingdom interfaces with rich, yet under-characterized spatial heterogeneity, where host and microbial metabolites can travel across the interface to exchange nutrients, induce responses, perform co-metabolism, etc^19,29–31^. Mapping the spatial distribution of metabolites across the host-microbe interfaces is thus critical to understand the underlying mechanisms of host-microbe interactions. Conventional metabolomics methodology involves liquid chromatography-mass spectrometry (LC-MS), for which luminal contents or fecal samples are usually separated from host tissues, resulting in the loss of spatial arrangement of host tissues and luminal contents. Mass spectrometry imaging (MSI) represents a powerful technique to visualize how molecules are spatially located within a sample and also quantify their abundances in these spaces^32,33^. Standard MSI sample preparation include freezing tissue samples and cryo-sectioning the intact frozen tissue, creating thin tissue slices with their chemical compounds and spatial arrangement preserved. The molecular contents of each spot across the sample surface are characterized, producing two-dimensional maps of relative analyte abundances across the sample^33^. Combined with other imaging modalities, such as histological staining^26,34^ and fluorescence in-situ hybridization ^35–37^, MSI can directly link tissue structures and cell types to specific detected molecules, offering unique perspectives into how detected chemical compounds are distributed across/within tissues and how functions might change due to disease states and environmental factors (i.e., chemical exposures, diet patterns, etc.).

We chose to study colon-lumen system, which represents an important site of host-microbe interactions. As the final part of the digestive system, the colon is the site for gut microbiota fermentation and harbors the greatest abundance of resident microbes in the gut^13,38^, making it a target to study host-microbe metabolism. Colonic is divided into three regions - proximal, mid, and distal- anatomical locations determined by the organization and consistency of luminal contents (i.e., fecal material within the colon prior to excretion)^39^. Continuous with the cecum, the proximal colon absorbs water and digestive nutrients within the luminal feces while mixing in mucus^39,40^. Next, the mid colon helps absorb remaining digestive nutrients, compacts luminal contents into a more solid state, and moves them along before reaching the distal colon, which enhances pressure to eliminate feces into the rectum^40^. Critically, the mid and distal colon also isolate the luminal contents from host tissue by forming a mucus barrier around the increasingly solid luminal contents^39^. Prior research identified associations between specific gut flora and colon diseases, such as colitis, Crohn’s disease, and colorectal cancer^41–45^. As the hotspot of microbial activity in the gut, the epithelium-lumen interface represents a way to directly map microbial effects on host tissue and potential health consequences^46,47^.

In this study, we aimed to systematically examine the proximal, mid, and distal regions of murine colons using MSI technology, aiming to reveal a comprehensive picture of metabolite spatial localization across the tissue-lumen interfaces (**Figure 1**). We collected proximal, mid, and distal colon tissues from male mice (*n*=4), using tissue samples containing luminal contents (i.e., feces) to preserve the tissue-lumen interfaces. We used matrix-assisted laser desorption/ionization (MALDI)-MS imaging, a popular ionization approach for MSI^33,48,49^ that uses chemical matrix to absorb laser energy and facilitate analyte ionization. MALDI-MS has been successfully applied to gut tissues^26,50–53^, while preserving the delicate tissue-lumen interface during sample preparation remains a challenge. We developed a “swiping” technique to successfully improve the preservation of luminal contents and tissue-lumen interfaces, particularly for the distal colon samples. In MSI experiments, we focused on characterizing the spatial distribution of molecules localized to different regions of these tissue: the host epithelium tissue itself, the luminal contents, and epithelium-lumen interface. MSI was done focusing on positively-charged ions with low molecular weight (<650 Da) and larger, negatively-charged compounds (i.e., lipids). We found 152 total features with clear spatial distribution patterns across the gut tissue-lumen interface in these joint analyses, highlighting potential microbial or microbially-modulated compounds as they are localized across the tissue-lumen interface and demonstrating robust and sensitive MALDI-MSI approaches for future investigations of gut microbe-host interactions.

**Figure 1.**
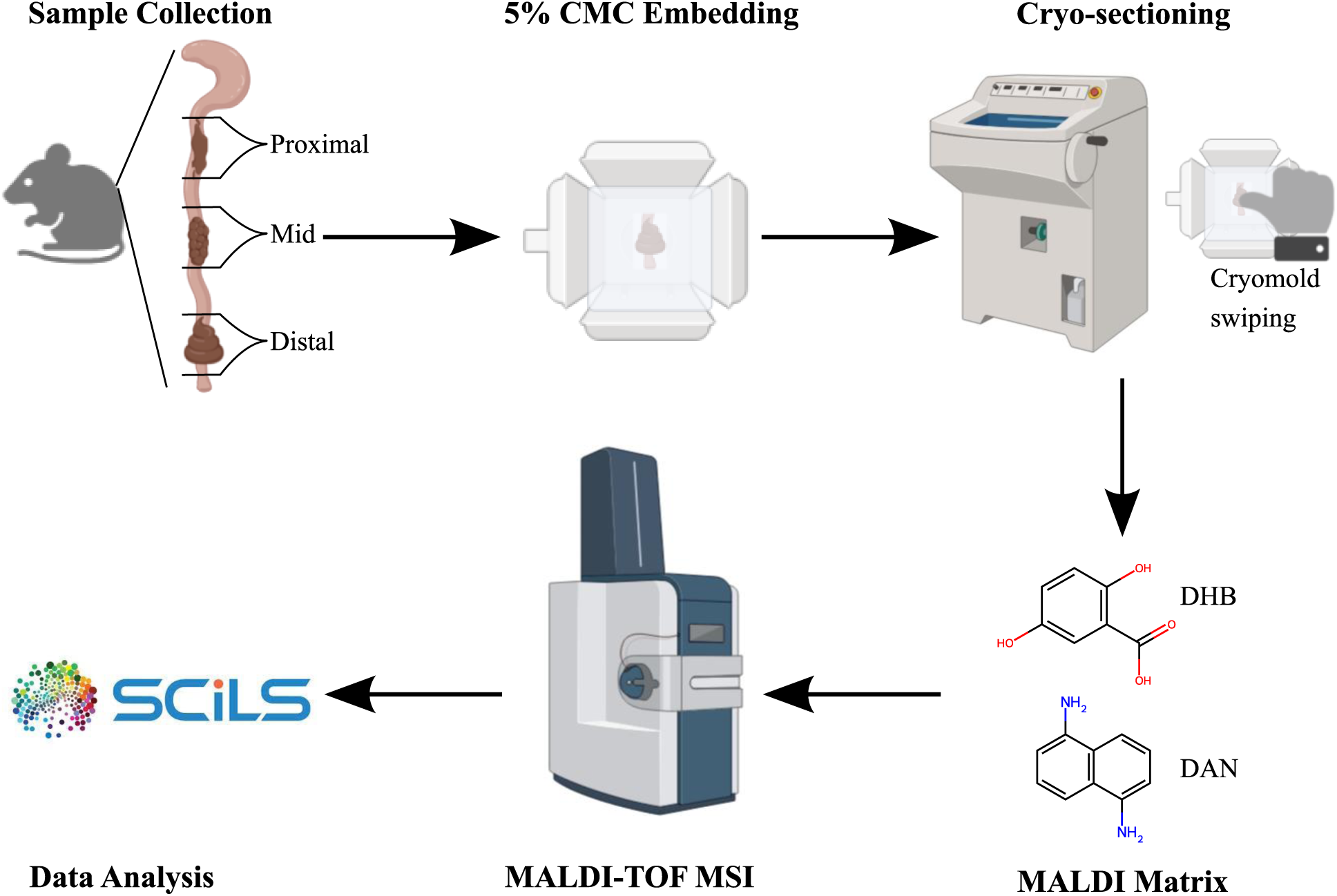
Experiment Design. Flow chart representing broad overview of experiment design, highlighting major steps. Images were produced by Biorender and edited in Inkscape. Proximal, mid, and distal colon tissues were harvested from four male mice. All tissues were embedded in 5% CMC and frozen via liquid nitrogen vapors before cryo-sectioning on a Leica cryostat. During sectioning, cryo-molds were briefly swiped with a gloved thumb for specific sections. Tissue sections were thaw-mounted onto ITO-coated slides and sprayed with DHB or DAN MALDI matrices. Matrix-sprayed slides underwent MALDI-TOF MSI on a Bruker timsTOF flex and the resulting data were analyzed using SCiLS Lab.

## Materials and Methods

### Sample Collection

A total of four wildtype C57BL/6 male mice (9 wks of age)) were collected for use in this project. For each use, the colon was set aside to separate into three regions: proximal, mid, and distal colon (**Figure 1**). Regions were identified by visual consistency of luminal contents and location/position on colon tissue. Generally, proximal colon was located just below cecum, mid colon was located around half-way down, and distal colon was anywhere in the lower third (excluding the final 1 cm, which was identified as the rectum and excluded). One tissue sample (approximately 0.5 – 1 cm length) was collected from each colonic region across all four mice, focusing on colonic areas containing feces. Three samples were collected from each of the four mice, totaling 12 samples. Each colon tissue sample was placed in a separate labeled 10x10x5 mm biopsy vinyl cryo-mold (Tissue-Tek). Molds were then filled with 5% carboxymethylcellulouse (CMC, viscosity 50-200 cP, c=4% H2O at 25°C; Cat# 419273, Sigma- Aldrich) embedding medium until tissue was completely immersed and cyromolds were filled.

CMC was chosen as an embedding medium over other tested media, such as M1 (Epredia) or 10% fish gelatin (Sigma-Aldrich), because CMC exhibited the best tissue preservation and fewest sectioning difficulties during early tests. Immediately after adding the 5% CMC solution, molds were slowly frozen via liquid nitrogen vapors to avoid introducing freeze-thaw artefacts and ice crystals^54–56^. Once molds were completely frozen, they were stored in dry ice during transfer to long-term storage at -80°C.

### Cryo-sectioning and the swiping technique

All cryo-sectioning was performed using a CM1850 Cryostat (Leica) at the MSU Center for Advanced Microscopy. For all cryo-molds, two separate indium tin oxide (ITO)-coated glass slides (Delta Technologies) were prepared for MALDI-MSI with different matrices. Each slide was cryo-sectioned to generate eight tissue cross-sections, each with a thickness of 16 μm at - 15°C. The eight tissue sections were organized into three groups representing cryo-sectioning swiping treatments. Serial tissue section collection for all slides occurred in the following order: sections 1-3 were not swiped prior to cryo-sectioning; sections 4-6 were all swiped briefly immediately before cryo-sectioning; and sections 7-8 were not swiped before cryo-sectioning (but were serial sections after sections 4-6 to examine if any swiping effects persist even without swiping). For swiping the respective sections, a gloved thumb was vertically moved/swiped twice across the cryo-mold before cryo-sectioning, with no added force or pressure. This swiping technique was particularly important for distal colon sections because the luminal contents are denser and less likely to lay flat during thaw-mounting. Cross-sectioning was chosen over transversal sectioning to enable better visualization of the host tissue interacting with fecal luminal contents. Each cross-section was immediately thaw-mounted onto the ITO-coated slide by holding a gloved finger or hand against the back of the slide until section was desiccated. One slide was prepared for each replicate and each colonic region, producing 24 total slides. This resulted in eight sections from each cryo-mold on each slide, with a total of 192 sections (*n*=96 for analysis with each MALDI matrix).

Once cryo-sectioning was concluded, slides were transferred on dry ice to the Qiu lab and subject to collection of bright-field optical images on a MZ10 F Modular Stereo Microscope (Leica) coupled with an Axiocam 202 Mono camera (Zeiss). Slides were then vacuum-dried and sealed using a vacuum sealer (FoodSaver) with sealing bags (Kirkland) and stored in -80°C until MALDI-MSI analysis.

### MALDI-MSI Sample Preparation

Prior to MALDI-MSI analysis, slides in vacuum packages were removed from -80°C and equilibrated at room temperature before opening to prevent condensation. Slides were placed in a vacuum chamber and desiccated using a PC 3012 NT Vario vacuum pump (Vacuubrand) connected to a nitrogen line for approximately 20 minutes. Following desiccation, slides were scanned using an Epson Perfection V850 Pro Scanner (Epson) to collect slide images for MSI.

### MALDI Matrix Application

All solvents used for MALDI matrix preparation and spraying were LC-MS grade. Analysis-ready sections were sprayed with 10 mg/mL of 2,5-dihydroxybenzoic acid (DHB; ThermoScientific; *n*=12) and 40 mg/mL 1,5-Diaminonaphthalene (DAN; TCI Chemicals; *n*=12), both in 70% acetonitrile (LC/MS Optima Fisher Chemical) and spiked with 0.1% trifluoroacetic acid (LC/MS Optima Fisher Chemical). Matrix spraying was done on a M3+ Sprayer (HTX Technologies). DHB spraying parameters were:: temperature: 75°C; pressure: 10 psi; flow rate: 100 μL/min; velocity: 1200 mm/min; track spacing: 2.5 mm; 4 total passes; CC pattern; no drying time; matrix density: 1.333 mg/mm^2^; and linear flow rate: 8.333e^-2^ mL/mm. DAN spraying parameters were: temperature: 75°C; pressure: 10 psi; flow rate: 120 μL/min; velocity: 1200 mm/min; track spacing: 2 mm; 4 total passes, CC pattern; no drying time; matrix density: 2 mg/mm^2^; and linear flow rate: 1e^-1^ mL/mm. Slides were stored in a dry nitrogen chamber until MALDI MSI analysis. Beyond matrix spraying, all slides were treated identically.

### MALDI-MSI

DHB and DAN samples were treated as separate batches for MSI analysis. All MSI was done on a timsTOF fleXtreme (Bruker Daltronics), a MALDI MS instrument with time-of-flight (TOF) mass analyzer, at 20 μm spatial resolution. For DHB sections, MSI was performed in positive ion mode and mass range of *m/z* 50-650. DHB laser parameters were: single beam; 1 burst of 200 shots; laser trigger frequency: 10,000 Hz; relative laser power: 65%; laser size: 20 μm x 20 μm; and raster width: 20 μm. Ion optics for DHB analysis were: MALDI plate offset: 50 V; deflection 1 delta: 60 V; funnel 1 RF: 120 Vpp; isCID energy: 0 eV; funnel 2 RF: 200 Vpp; multipole RF: 200 Vpp; quadrupole ion energy: 5 eV; low mass: 50 *m/z*; focus pre-TOF transfer time: 55 μs; and pre-pulse storage: 3 μs. For DAN analysis, MSI was performed in negative ion mode with a mass range of *m/z* 100-1100 to enable coverage of negative ions and lipids. DAN laser parameters were: single beam; 1 burst of 200 shots; laser trigger frequency: 10,000 Hz; relative laser power: 60%; laser size: 20 μm x 20 μm; and raster width: 20 μm. Ion optics for DAN analysis were: MALDI plate offset: 50 V; deflection 1 delta: -70 V; funnel 1 RF: 300 Vpp; isCID energy: 0; funnel 2 RF: 200 Vpp; multipole RF: 200 Vpp; quadrupole ion energy: 5 eV; low mass: 100 *m/z*; focus pre-TOF transfer time: 65 μm; and pre-pulse storage: 8 μs.

### Histological Staining

After MSI analysis, slides were briefly plunged (4 seconds max) into LC-MS acetonitrile (LC/MS Optima Fisher Chemical) to rinse off MALDI matrices for hematoxylin and eosin (H&E) histological staining. H&E staining was done by: rehydration with 100% ethanol for 40 s, transfer to new 100% ethanol for 40 s, 95% ethanol for 40 s, 80% ethanol for 40s, and water for 1 min; staining: hematoxylin 560 (Leica; catalog: 3801576) for 3 mins; differentiation: water for 1 min, Define (Leica) for 20 s, water for 1 min, blue buffer 8 (Leica) for 40 s, water for 1 min, and 95% ethanol for 40 s; counterstain: alcoholic eosin Y515 (Leica) for 12 s; dehydration: 95% ethanol for 40 s, transfer to new 95% ethanol for 40 s, 100% ethanol for 2 min, transfer to new 100% ethanol for 2 min, and transfer to new 100% ethanol for 3 min; and clearing: Histo-Clear (National Diagnostics) for 3 min, and transfer to new Histo-Clear for 3 mins^57^. After staining, slides were mounted with a plastic cover slip. H&E-stained bright-field images were acquired on an Axioscan 7 Microscope Slide Scanner (Zeiss).

### Data Analysis and Processing

MALDI-MSI data for both DHB and DAN analyses were imported into SCiLS Lab v2024b Pro (Bruker Daltronics). Importing parameters were: bin size: 1.5 mDa; at *m/z*: 300; and number of bins: 696250. Feature finding parameters were: TIC normalization; T-ReX^2^ algorithm; spatial noise filtering: weak; 10% coverage; intensity threshold: 0.5% relative intensity; and auto *m/z* interval width. Localized ions were visually identified by examining features passed the filtering criteria. Ions were grouped into one of eight localization categories based on visual inspections for their locations of highest spatial relative abundance. These categories were: lumen (localized exclusively to the luminal contents); lumen and interface (localized within the luminal contents and extending outside lumen); tissue inner layer (localized within tissue but not present throughout tissue, lumen, or interface); tissue all (localized across all tissue, with generally uniform abundance throughout); tissue outer layer (localized to outermost tissue edges); tissue and lumen (localized in luminal contents and tissue areas); interface enriched (localized primarily to interface region, but otherwise absent in lumen and tissue); and absent (referring to ions not detected in a specific section but present in other sections). Image modifications and edits were done using Biorender and Inkscape v1.2^58^.

## Results and Discussion

### Swiping during cryo-sectioning improves sample integrity without delocalizing ions

Due to the dry, solid nature of luminal contents in the distal colon, we noticed that dense components of the luminal contents do not readily lay flat during thaw-mounting. This feature biases MSI analysis as the MALDI laser will not properly focus on sample surface and ionize analytes where tissue surface is not flat; no data will be acquired in those areas. One simple workaround involves swiping a gloved finger across the cryo-mold prior to sectioning, which yields more intact sections with flatter luminal contents. This technique was adopted during our testing and improved the integrity of colon sections (**Figure 2; Tables S1-S2**). The greatest improvements from swiping were observed in distal colon sections, where all tissue and lumen integrities were significantly improved as a result of swiping (**Figure 2; Tables S1-S2**). Mid colon samples displayed marginal benefits from swiping, with some samples exhibiting much better tissue/lumen integrity from swiping while others were marginal improvements. For proximal colon samples, there were no visible benefits from swiping as tissue and lumen integrities were nearly-identical across all samples (**Tables S1-S2**). Since the swiping technique offered great improvements to sample integrity for some samples and marginal for others without major adverse effects, we continued using this technique throughout cryo-sectioning.

**Figure 2.**
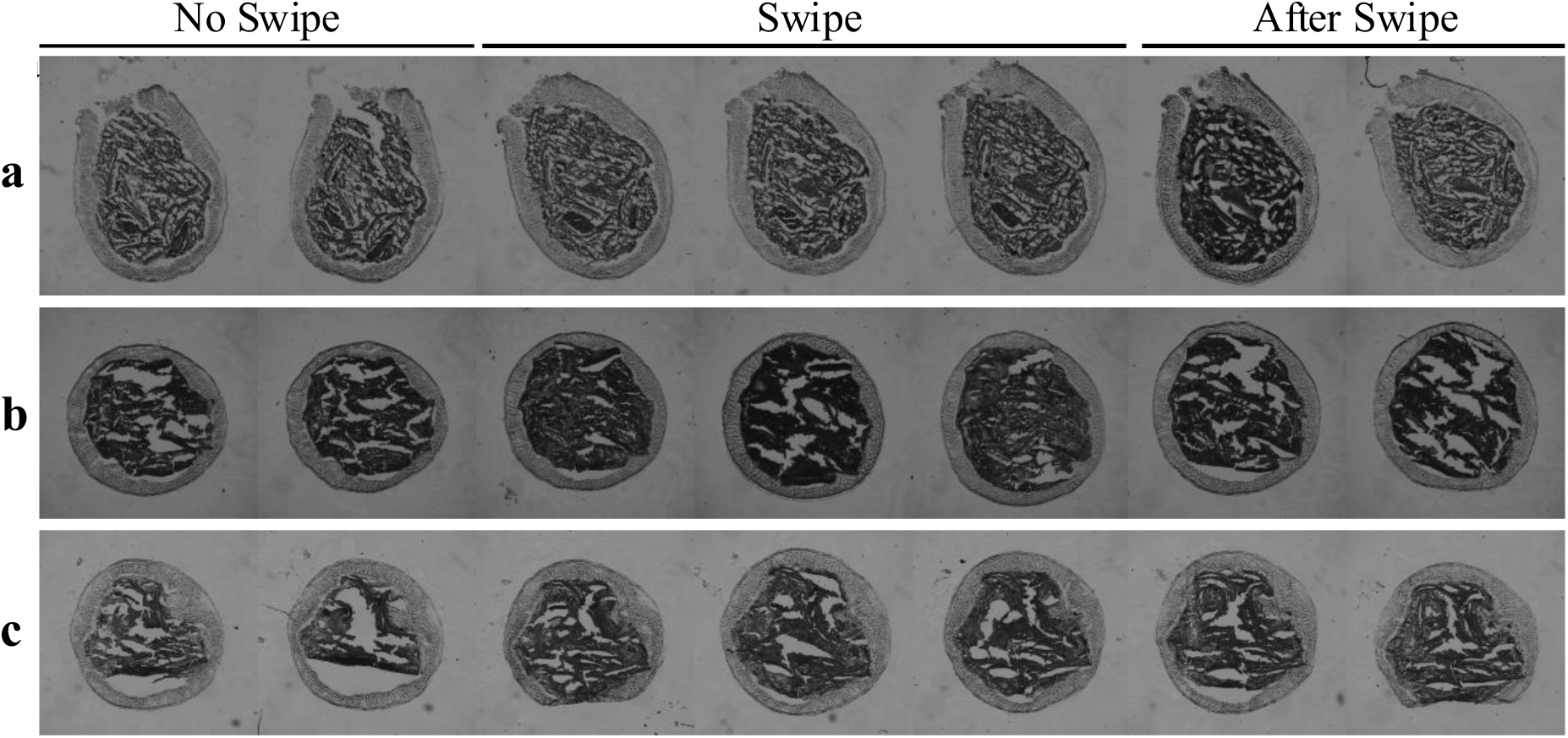
Swiping before cryo-sectioning improves sample integrity. Optical bright-field images demonstrate swiping prior to cryo-sectioning results in more intact tissue and luminal contents across serial sections. **a**, Proximal colon, mouse replicate three:; **b**, Mid colon, mouse replicate three; **c**, Distal colon, mouse replicate one.

We recognized that while sample integrity was improved, this technique can potentially cause delocalization of molecules, particularly small molecules; however,we did not observe signs of delocalization effects from swiping across the tested samples and MALDI matrices (**Figure 3**). To test potential analyte delocalization from swiping, we performed serial sections where the first three acted as non-swiped controls, followed by three sections swiped right before sectioning and three further serial sections without swiping the tissue to act as post-swiping controls. These serial sections were mounted onto the same slide for back-to-back comparisons in MSI experiments. **Figure 3** shows that ions localized to the luminal contents, epithelium tissue, and epithelium-lumen interface were still localized to their respective regions after swiping. By exposing the sample to a swift swiping of a gloved finger, the CMC-embedded cryo-mold is theoretically momentarily melted, which is expected to result in delocalization of molecules in tissues and creating a “smear” following swiping direction. However, there were neither such smearing patterns nor ion delocalization in the MSI data (**Figure 3**), suggesting this brief heating and flattening does not compromise tissue integrity nor molecules’ spatial distribution in the cryo-mold. We noticed that putting thumbs against the mold without moving caused severe ion delocalization; thus, the swift action of swiping seems to be a key for the success of the technique. We postulate that during the swift swiping movement, the cryo-mold was momentarily heated and very quickly re-frozen as the finger leaves due to the -20C internal temperature of the instrument, avoiding the diffusion of molecules on surface, while the swiping movement creates a thin layer of embedding media to help with section integrity. Further work is desired to explore the broad applicability of this swiping technique in other sample types and embedding media, while our results illustrate swiping during cryo-sectioning maintains colon tissue morphology without delocalizing detected ions in 5% CMC. Thus, we proceeded with the swiping method going forward.

**Figure 3.**
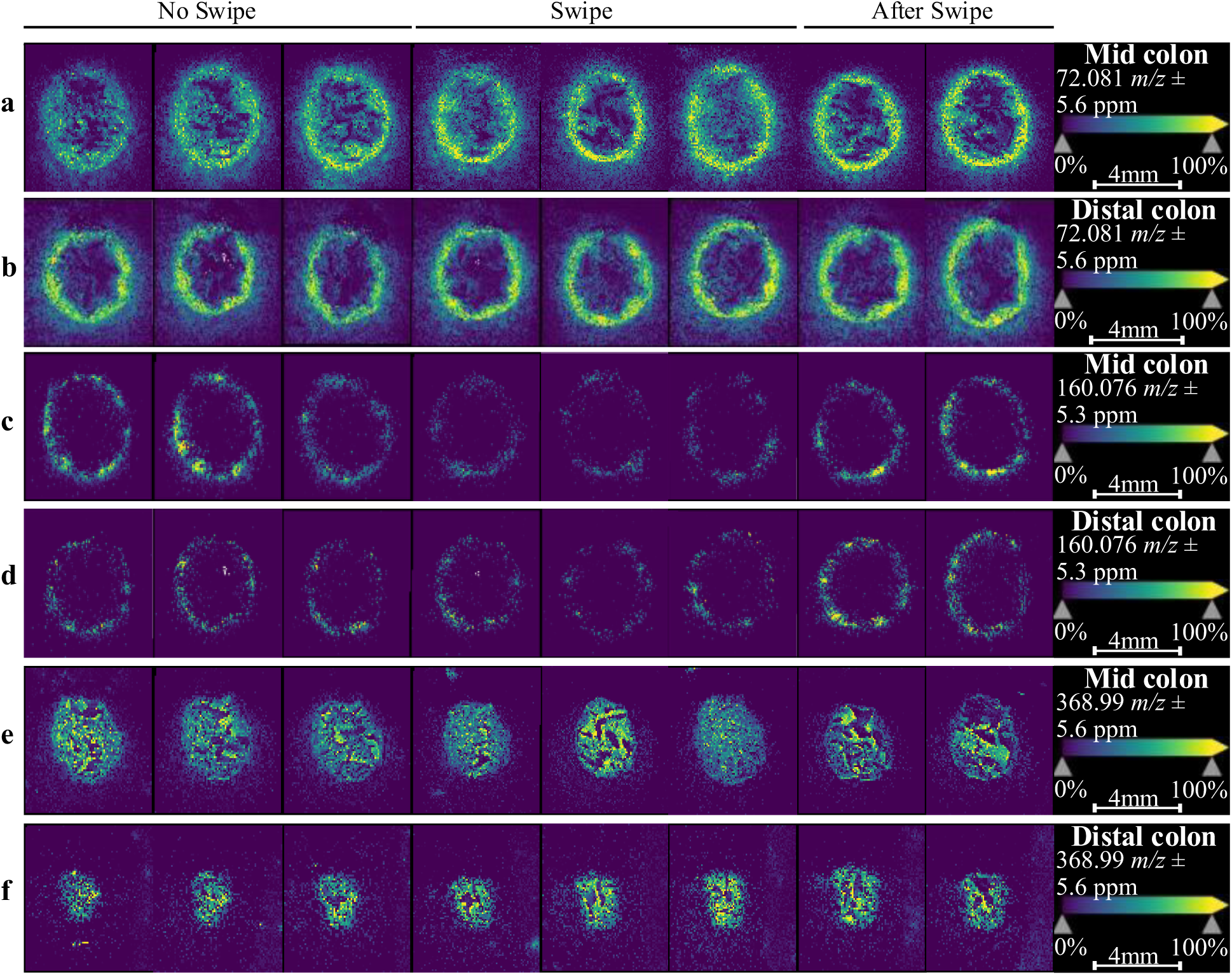
Swiping during cryo-sectioning does not affect ion localization. All sub-figures are DHB-sprayed slides. Individual ions are represented in each heatmap sub- figure, where ion abundance is represented by bright (yellow) colors and ion absence is visualized by negative (purple) space. All slides have eight sections in the following order: Sections 1-3, no swipe; Sections 4-6, swiped; Sections 7-8, no swipe (serial sections after swiping to examine if swiping effects persist). Representative slides and ions show there are no obvious delocalization effects caused by swiping the cryo-mold during cryo-sectioning. **a**, Mid colon sections from mouse two displaying ion localized to tissue and lumen (*m/z* 72.0806). **b**, Distal colon sections from mouse four displaying ion localized to tissue (*m/z* 72.0806). **c**, Mid colon sections from mouse four displaying ion localized to tissue inner layer (*m/z* 160.0758). **d**, Distal colon sections from mouse four displaying ion localized to tissue inner layer (*m/z* 160.0758). **e**, Mid colon sections from mouse three displaying ion localized to luminal contents (*m/z* 368.9892). **f**, Distal colon sections from mouse two displaying ion localized to luminal contents (*m/z* 368.9892).

The swiping technique greatly improved the cryo-sectioning of colon-lumen samples, while challenge persists to acquire perfectly flat luminal contents after vacuum desiccation of the mounted sections (**Figure 2**; **Tables S1-2**). Namely, the vacuum desiccation and matrix spraying both present opportunities for luminal contents (especially in distal colon) to completely dry out and crack, dislodging themselves from the slide surface. This can be partly mitigated by using spraying parameters creating a ‘wet’ spray to weigh down the luminal contents; drier sprays often worsened sample integrity during our tests. Other alternative solutions include coating slides with poly-L-lysine, such as used by Kamphius and colleagues^39^. We tested poly-L-lysine coatings during method optimization and did not observe notable benefits. Meanwhile, the swiping technique consistently improved sample integrity, with or without poly-L-lysine coatings. Another method for improving sample integrity during sectioning involves adhesive tapes, like those used by Bender and colleagues^59^, to help tissue lay flat. However, these tapes can be expensive and potentially cause loss of luminal contents (especially in distal colon where luminal contents can be easily dislodged). Ultimately, more work is needed to investigate the underlying mechanisms affecting our observations from swiping, but the swiping technique employed here helps maintain the spatial and histological integrity of colon tissues without observable analyte delocalization.

### Colonic regions exhibit distinct molecular spatial heterogeneity

Detected ions were consistently localized to specific areas of the colon sections, across different colonic regions (**Figures 4-5**). A total of 51 and 101 ions were localized to the granular eight localization categories in DHB and DAN, respectively, across different colonic regions and samples. For DHB-sprayed samples analyzed in a low-molecular weight metabolite range (positive mode, 50-650 *m/z*), examples include *m/z* 71.0728 and *m/z* 296.0659 for tissue, and *m/z* 192.981 and *m/z* 365.0178 for lumen (**Figure 4; Table S3**). Other granular identifications for DHB slides are *m/z* 146.9818 and *m/z* 204.123 for tissue outer layer, *m/z* 180.0655 for tissue inner layer, and *m/z* 184.073 and *m/z* 496.3394 for both tissue and lumen (**Table S3**). Such assignments indicate these ions’ spatial distribution is maintained across the colon. For DAN- sprayed samples analyzed in a larger metabolite/lipid range (negative mode, 100-1100 *m/z*), a total of 101 ions exhibited discrete spatial distributions, presenting more granularity in spatial distribution patterns compared to the DHB-sprayed samples (**Figure 5; Table S4**). Examples included *m/z* 214.0482 and *m/z* 391.2258 localizing to whole tissue across all regions, *m/z* 471.2415 and *m/z* 796.5367 localizing to the luminal contents, *m/z* 863.564 localizing to inner tissue, and *m/z* 794.5711 localizing to outer tissue (**Figure 5; Table S4**). Some ions were localized to the boundaries between the tissue and lumen, such as *m/z* 152.9957 and *m/z* 255.2327 localizing to both tissue and lumen, or *m/z* 465.3046 localizing between the luminal contents and interface site (**Figure 5; Table S4**). We notice that localization assignments frequently varied between regions. For example, in the DHB dataset, *m/z* 129.1385 localized to inner tissue layer for proximal/mid colon but was tissue/lumen for distal colon, or *m/z* 250.0935 assigned as inner tissue layer for proximal, all tissue for mid colon, and lumen for distal colon (**Table S3**). Some varyingly-localized DAN ions include *m/z* 1021.522 as whole tissue-localized in proximal colon to outer tissue-localized for mid and distal colon, or *m/z* 161.0458 localized to tissue/lumen for proximal and mid colon while enriched at the interface for distal colon (**Table S4**). These patterns likely reflect the varying functions of the distinct colonic regions, which are likely driven by the combination of gut tissues physiology, food digestion and microbiota heterogeneity along the colonic region.

**Figure 4.**
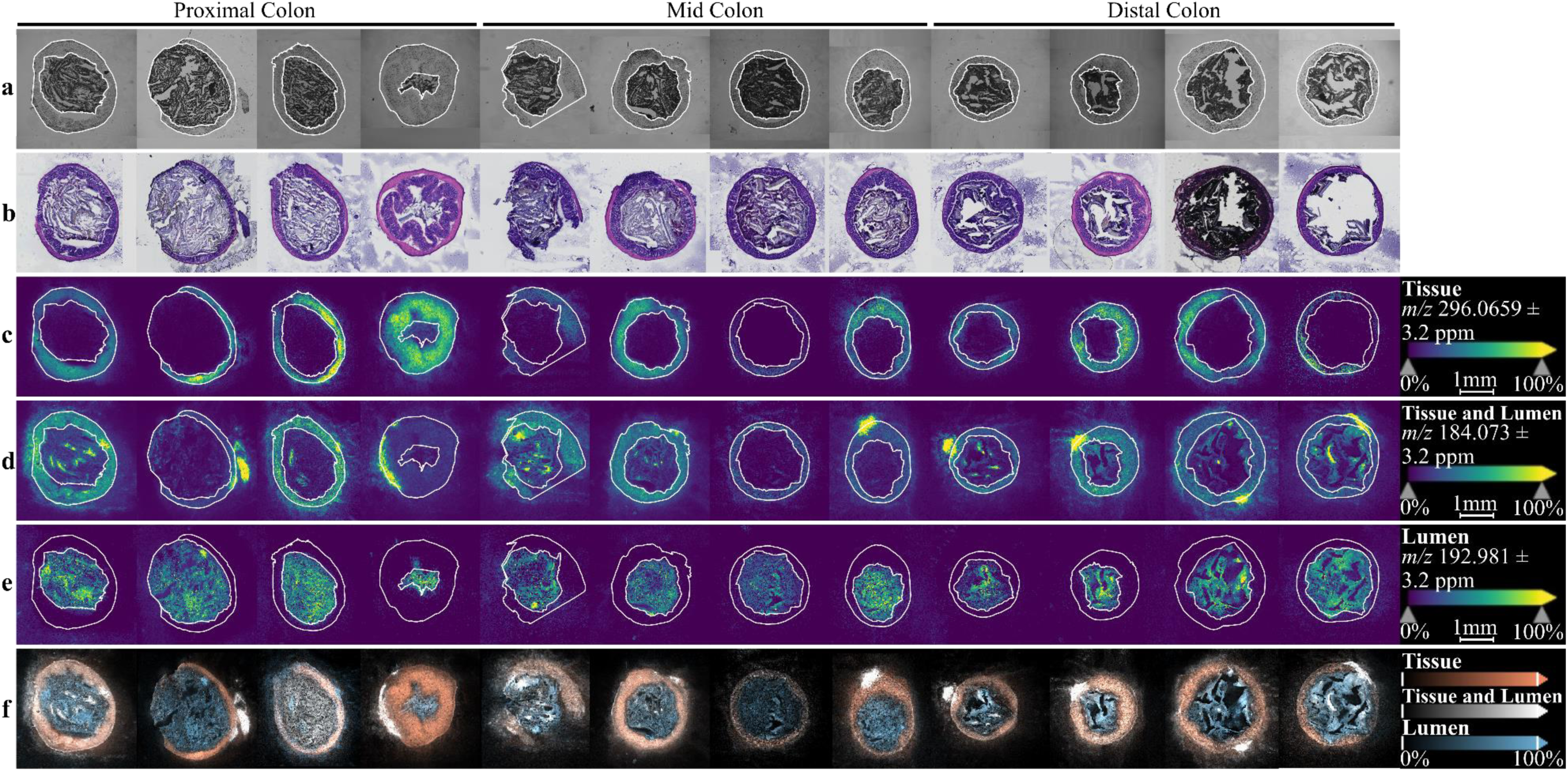
Colon tissue exhibits tissue and lumen molecular spatial heterogeneity across colonic regions for DHB-sprayed sections. All sub-figures are DHB-sprayed slides analyzed. Individual ions are represented in each heatmap sub-figure, where ion abundance is represented by bright colors and ion absence is visualized by negative space. All rows represent a single representative section from the same animal replicate, while each column displays the same section for all animal replicates grouped by colonic region White lines in **a-e** are tracings around the outer tissue edges and the interface, to help visualize discrete boundaries between the lumen and tissue. **a**, Representative bright-field images. **b**, Representative H&E-stained image for each section. **c**, Ion localized to tissue (*m/z* 296.0659). **d**, Ion localized to tissue and lumen (*m/z* 184.073). **e**, Ion localized to luminal contents (*m/z* 192.871). **f**, Ions from **c-e** overlaid together and color-coded according to localization category.

**Figure 5.**
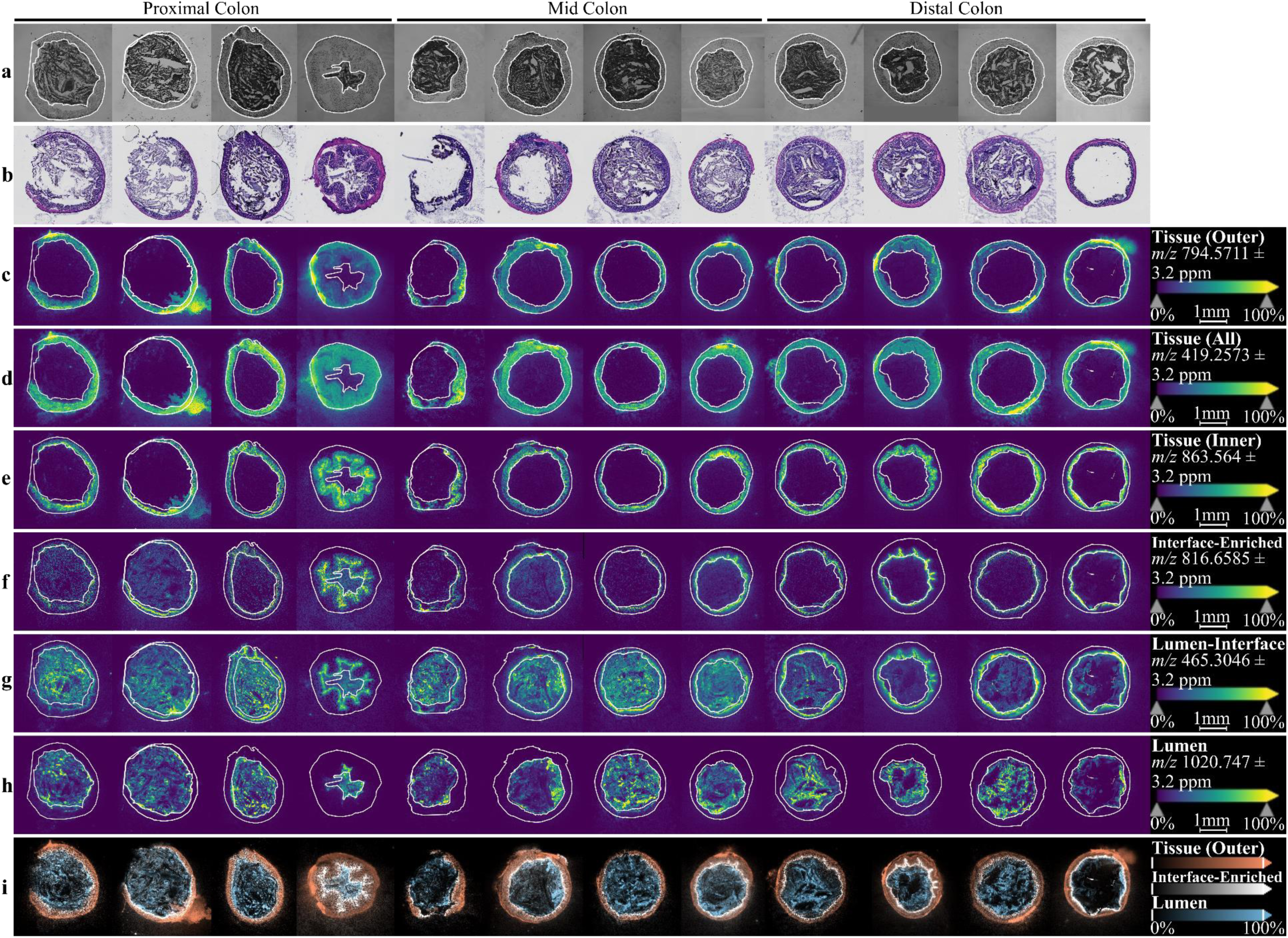
DAN MALDI matrix-sprayed slides demonstrate robust colonic molecular spatial heterogeneity. All sub-figures are DAN-sprayed slides analyzed. Individual ions are represented in each heatmap sub-figure, where ion abundance is represented by bright colors and ion absence is visualized by negative space. All rows represent a single representative section from the same animal replicate, while each column displays the same section for all animal replicates grouped by colonic region.. White lines in **a-h** represent tracings around the outer tissue edge and the interface regions, to help visualize the discrete boundaries between the lumen and tissue. **a**, Representative bright-field images. **b**, Representative H&E-stained image for each section. **c**, Ion localized to outer tissue layer (*m/z* 794.5711). **d**, Ion localized to all tissue area (*m/z* 419.2573). **e**, Ion localized to inner tissue layer (*m/z* 863.564). **f**, Ion enriched in interface area (*m/z* 816.6585). **g**, Ion localized to lumen and interface areas (*m/z* 465.3046). **h**, Ion localized to luminal contents (*m/z* 1020.747). **i**, Ions from **c**, **f**, and **h** overlaid together and color-coded according to localization category.

Amongst all spatial localization patterns, we are particularly interested in the interface, where the gut tissue and luminal contents meet in proximity to each other. We found a list of interface-enriched ions in DAN-sprayed samples while none was found in DHB-sprayed samples. Examples of these DAN interface ions include *m/z* 506.2887, *m/z* 728.5591, *m/z* 726.5445, *m/z* 154.0516, *m/z* 788.5546, and more (**Figures 5-6**; **Table S4**). Interestingly, many ions whose localization changed between colonic regions were tissue and/or lumen-localized for proximal/mid colon but were interface-enriched for distal colon. Examples include *m/z* 844.6902, *m/z* 478.2952, *m/z* 244.0832, and *m/z* 837.5503 (**Table S4**). This may be related to the differences of mucus layers in different regions. In the proximal colon, there is no sterile mucus layer separating host tissue from bacteria-laden luminal contents, while the luminal contents desiccate and compact traveling down the mid and distal colon with more defined mucus barrier between luminal contents and host tissues.^39^ Some other interface-enriched ions were shared across the colonic regions, suggesting that they may have consistent roles. Very interestingly, we observed different patterns for interface-enriched ions, where some ions seem to localize right along the luminal contents boundary like *m/z* 154.0516 (**Figure 6a**) while others localize along the epithelial tissue like *m/z* 816.6585 (**Figure 5f**), *m/z* 506.2887 (**Figure 6b**), or *m/z* 788.5546 (**Figure 6c**). These distinct inter-interface localizations hint at discrete molecular activities occurring within the interface site. For example, the along-lumen interface ions potentially reflect components of the mucus barriers that originated from luminal contents, either food or microbial activities, and the along-tissue interface ions highlight the mucosa tissue layers. While these ions are all enriched in the tissue-lumen interface site, their detailed spatial localization reveals their potential sources and physiological roles at the gut-lumen interface. Future analyses can help us to detect, identify, and investigate these analytes according to Metabolomics Standards Initiative (MSI) guidelines^60^ and understand the biological significance of these interface-enriched metabolites.

**Figure 6.**
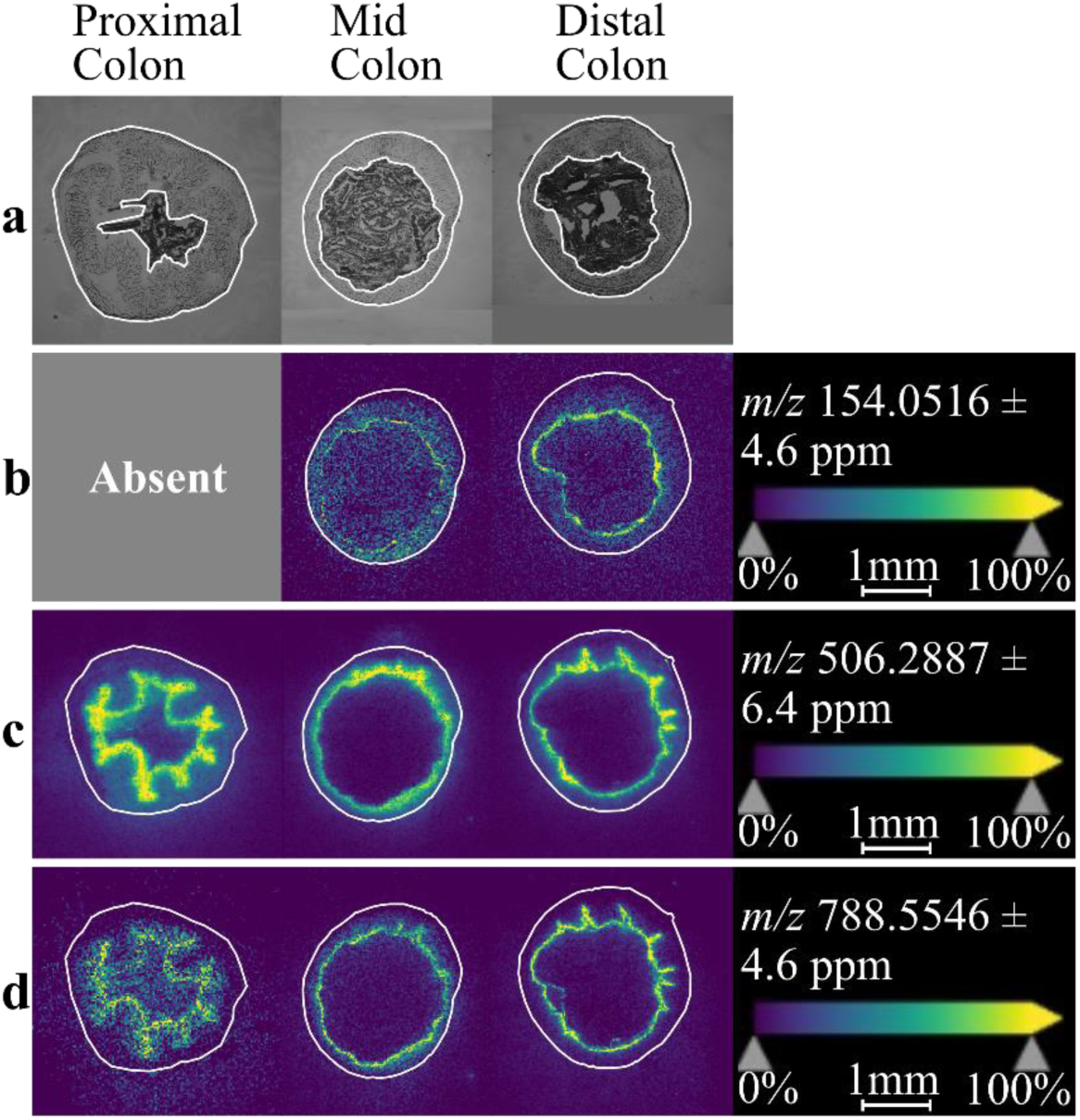
Interface-enriched ions are detectable for DAN-sprayed sections across all colonic regions. All slides were sprayed with DAN. A single representative interface-enriched ion is displayed for each colonic region. Each row contains the same animal replicate with a single section from each colonic region (columns). Representative proximal, mid, and distal colon samples were from animals four, four, and two, respectively. **a**, Bright-field images. White lines are tracings around the tissue and interface regions for each sections. For **b-d**, only tracings around the tissue are visualized to avoid overshadowing the interface region in each ion image. **b**, Interface-enriched ion *m/z* 154.0516 (absent in all proximal colon samples). **c**, Interface-enriched ion *m/z* 506.2887. **d**, Interface-enriched ion *m/z* 788.5546.

### Histological Analyses

Post-MALDI hematoxylin and eosin (H&E) staining of colonic tissue offered insights into tissue morphology and preservation across sample treatment during this project. These histological stains also enable greater visualization of the epithelium-lumen interface, by inspecting the negative space between the luminal contents and host tissue. Our H&E-stained images demonstrate good tissue integrity, although luminal contents were not always preserved.

For proximal and mid colon samples, the epithelial tissue and luminal contents were maintained, but luminal contents were consistently absent or depleted in distal colon sections (**Figures 4-5**). The absence of luminal contents in distal colon sections is likely the result of desiccated luminal contents being dislodged during the post-MALDI washing process and H&E staining procedure. This pattern highlights the crucial role sample preparation and treatment plays during histological and molecular analyses. In general, these H&E-stained images depict the expected morphology of the proximal, mid, and distal colon tissues. For example, the larger, horizontal-to-vertical orientation of intestinal villi (**Figure 4b**; **Figure 5b**) is expected in proximal colon because it deals with looser, more liquid luminal contents and is focused on recovering critical nutrients, removing and compacting undigested food products, and extracting water. Meanwhile, mid and distal colon focus on pressurizing and compacting the luminal contents, so their tissue walls are more organized vertically, with smaller villi and crypts (**Figure 4b**; **Figure 5b**). Such patterns reinforce similar findings from previous literature^39^. Together, these images demonstrate the morphological differences across the colon (**Figure 4b**; **Figure 5b**), which are reflected in the molecular spatial heterogeneity of the colon (**Figures 4-5**).

One possible explanation for the shifting distribution of interface-localized ions between the colonic regions is that these interface ions actually represent mucus layer ions. These ions still localize to the interface site, but this also is where the mucus layers can form. As these mucus layers shift and become incorporated into the luminal contents while new layers envelop the formed fecal pellet, these ions will move around the space accordingly. As we demonstrated the successful post-MALDI H&E staining for gut-lumen tissue samples, future work can apply mucin-specific staining methods to visualize the spatial localization of mucus layers and investigate their spatial co-localization with interface and lumen ions.

## Conclusions and Future Directions

In this project, we developed a MALDI-MSI workflow to map the metabolomic landscape across the colon epithelium-lumen interface, the hotspot for host-microbe interactions, at multiple colonic regions. With a swiping technique at cryo-sectioning, we successfully improved the preservation of gut luminal contents and demonstrated that the swift swiping did not cause delocalization of metabolites on tissue. We found a total of 162 features with clear spatial localization patterns in gut-lumen cross sections from MALDI-MSI analysis with two different matrices for positive and negative modes; the spatial distribution patterns ranges from tissue only (outer tissue, inner tissue and all across tissue), lumen only, lumen and tissue and gut-lumen interface-enriched. The distribution of these ions also varies between colonic regions, which may reflect distinct molecular mechanisms associated with microbial activity in these areas. Overall, our results revealed unprecedented details regarding the spatial metabolomic landscape across the colon-lumen interfaces that is closely related to the spatial arrangement of tissue structure, mucus layer and luminal contents. Further work will focus on annotation and structural elucidation of key species with spatial localizations relevant to gut-microbiome interactions, developing data analysis pipelines for mining the imaging dataset, further expanding the coverage of metabolites with other MALDI matrices, and coupling the MALDI-MSI with other imaging modalities. Furthermore, the methods and analyses presented here demonstrate the greater need for considering spatial distribution when investigating host-microbiome-metabolome relationships and provide new directions for exploring the spatiality of the gut microbiome and host health.

## Acknowledgements

The authors thank our collaborators Alexandra Chapman, Jaewon Sim, and Dr. Geoffroy Laumet for contributing samples and assistance with sample collection for this project. We also thank Dr. Melinda Frame for use of the Center for Advanced Microscopy’s cryostat and for her training on the instrument. We would also like to thank Amy Porter at MSU’s Histopathology Lab for her guidance and suggestions during early stages of project planning. We are also grateful to Dr. Keith MacRenaris and Dr. Aaron Sue at MSU’s Quantitative Bio-Element Analysis and Mapping Center for training and use of the facility’s slide scanner. Lastly, we thank Dr. Tony Schilmiller and the MSU Mass Spectrometry and Metabolomics Core for use of their facility’s HTX M3+ Sprayer.

## Author Contributions

T.A.Q. conceived and designed the project with J.J.H. Sample collection was done by J.J.H. and S.H.A. under direction from T.A.Q. J.J.H. performed all experimental lab work, collected MSI data, and performed data analysis with guidance from T.A.Q. S.H.A performed H&E staining with J.J.H. assisting. J.J.H. wrote the manuscript with contributions from T.A.Q. All authors reviewed the final manuscript.

## Ethics Approval

Animal experiments were performed following the animal use protocol PROTO202300090 approved by the Institutional Animal Care and Use Committee (IACUC) at Michigan State University.

## References

1. Thursby, E. & Juge, N. Introduction to the human gut microbiota. Biochem. J. 474, 1823– 1836 (2017).

2. Natividad, J. M. M. & Verdu, E. F. Modulation of intestinal barrier by intestinal microbiota: Pathological and therapeutic implications. Pharmacol. Res. 69, 42–51 (2013).

3. Bosco, N. & Noti, M. The aging gut microbiome and its impact on host immunity. Genes Immun. 22, 289–303 (2021).

4. Gensollen, T. et al. How colonization by microbiota in early life shapes the immune system. Science. 2016;352(6285):539-544. doi:10.1126/science.aad9378. *Science* **352**, 539–544 (2016).

5. Corbin, K. D. et al. Host-diet-gut microbiome interactions influence human energy balance: a randomized clinical trial. Nat. Commun. 14, (2023).

6. Murphy, E. F. et al. Composition and energy harvesting capacity of the gut microbiota: Relationship to diet, obesity and time in mouse models. Gut 59, 1635–1642 (2010).

7. Yadav, H. Gut Microbiome Derived Metabolites to Regulate Energy Homeostasis: How Microbiome Talks to Host. J. Postgenomics Drug Biomark. Dev. 6, (2016).

8. van Schaik, W. The human gut resistome. Philos. Trans. R. Soc. B Biol. Sci. 370, (2015).

9. Yoon, M. Y., Lee, K. & Yoon, S. S. Protective role of gut commensal microbes against intestinal infections. J. Microbiol. 52, 983–989 (2014).

10. Banerjee, P. et al. Digestion and gut microbiome. in *Nutrition and Functional Foods in Boosting Digestion*, Metabolism and Immune Health 123–140 (Academic Press, 2021). doi:10.1016/B978-0-12-821232-5.00029-X

11. Bezirtzoglou, E. et al. Maintaining digestive health in diabetes: The role of the gut microbiome and the challenge of functional foods. Microorganisms 9, 1–26 (2021).

12. Bull, M. J. & Plummer, N. T. Part 1: The Human Gut Microbiome in Health and Disease. Metagenomics 197–213 (2014). doi:10.1016/B978-0-08-102268-9.00010-0

13. De Vos, W. M., Tilg, H., Van Hul, M. & Cani, P. D. Gut microbiome and health: mechanistic insights. Gut 71, 1020–1032 (2022).

14. Murugesan, S. et al. Gut microbiome production of short-chain fatty acids and obesity in children. Eur. J. Clin. Microbiol. Infect. Dis. 37, 621–625 (2018).

15. Ross, F. C. et al. The interplay between diet and the gut microbiome: implications for health and disease. Nat. Rev. Microbiol. (2024). doi:10.1038/s41579-024-01068-4

16. Johnson, C. H., Ivanisevic, J. & Siuzdak, G. Metabolomics: Beyond biomarkers and towards mechanisms. Nat. Rev. Mol. Cell Biol. 17, 451–459 (2016).

17. Patti, G. J., Yanes, O. & Siuzdak, G. Metabolomics: the apogee of the omic trilogy. Nat Rev Mol Cell Biol 13, 263–269 (2012).

18. Viant, M. R., Kurland, I. J., Jones, M. R. & Dunn, W. B. How close are we to complete annotation of metabolomes? Curr. Opin. Chem. Biol. 36, 64–69 (2017).

19. Collins, S. L., Stine, J. G., Bisanz, J. E., Okafor, C. D. & Patterson, A. D. Bile acids and the gut microbiota: metabolic interactions and impacts on disease. Nat. Rev. Microbiol. 21, 236–247 (2023).

20. Fontdevila, L. et al. Examining the complex Interplay between gut microbiota abundance and short-chain fatty acid production in amyotrophic lateral sclerosis patients shortly after onset of disease. Sci. Rep. 14, 1–10 (2024).

21. Santoru, M. L. et al. Cross sectional evaluation of the gut-microbiome metabolome axis in an Italian cohort of IBD patients. Sci. Rep. 7, 1–14 (2017).

22. Yang, W. et al. Intestinal microbiota-derived short-chain fatty acids regulation of immune cell IL-22 production and gut immunity. Nat. Commun. 11, 1–18 (2020).

23. Zierer, J. et al. The fecal metabolome as a functional readout of the gut microbiome. Nat. Genet. 50, 790–795 (2018).

24. Ahmed, I., Roy, B. C., Khan, S. A., Septer, S. & Umar, S. Microbiome, metabolome and inflammatory bowel disease. Microorganisms 4, 1–19 (2016).

25. Wu, R., Xiong, R., Li, Y., Chen, J. & Yan, R. Gut microbiome, metabolome, host immunity associated with inflammatory bowel disease and intervention of fecal microbiota transplantation. J. Autoimmun. 141, 103062 (2023).

26. Ponzoni, A. et al. An untargeted metabolomic study using MALDI-mass spectrometry imaging reveals region-specific biomarkers associated with bowel inflammation. Metabolomics 21, (2025).

27. Spruill, M. L., Maletic-Savatic, M., Martin, H., Li, F. & Liu, X. Spatial analysis of drug absorption, distribution, metabolism, and toxicology using mass spectrometry imaging. Biochem. Pharmacol. 201, 832–842 (2022).

28. Tropini, C., Earle, K. A., Huang, K. C. & Sonnenburg, J. L. The Gut Microbiome: Connecting Spatial Organization to Function. Cell Host Microbe 21, 433–442 (2017).

29. Gao, K., Mu, C. L., Farzi, A. & Zhu, W. Y. Tryptophan Metabolism: A Link between the Gut Microbiota and Brain. Adv. Nutr. 11, 709–723 (2020).

30. Li, T. T. et al. Microbiota metabolism of intestinal amino acids impacts host nutrient homeostasis and physiology. Cell Host Microbe 32, 661–675.e10 (2024).

31. Shan, Y., Lee, M. & Chang, E. B. The Gut Microbiome and Inflammatory Bowel Diseases. Annu. Rev. Med. 73, 455–468 (2022).

32. Cole, L. M. Imaging Mass Spectrometry: Methods and Protocols. (Humana Press, 2017).

33. Greaves, J. & Roboz, J. Mass Spectrometry for the Novice. (CRC Press, 2014).

34. Bai, H., Linder, K. E. & Muddiman, D. C. Three-dimensional (3D) imaging of lipids in skin tissues with infrared matrix-assisted laser desorption electrospray ionization (MALDESI) mass spectrometry. Anal. Bioanal. Chem. 413, 2793–2801 (2021).

35. Bourceau, P. et al. Visualization of metabolites and microbes at high spatial resolution using MALDI mass spectrometry imaging and in situ fluorescence labeling. Nat. Protoc. 1–30 (2023). doi:10.1038/s41596-023-00864-1

36. Geier, B. et al. Spatial metabolomics of in situ host–microbe interactions at the micrometre scale. Nat. Microbiol. 5, 498–510 (2020).

37. Huber, K. et al. Multimodal analysis of formalin-fixed and paraffin-embedded tissue by MALDI imaging and fluorescence in situ hybridization for combined genetic and metabolic analysis. Lab. Investig. 99, 1535–1546 (2019).

38. Guarner, F. & Malagelada, J.-R. Gut flora in health and disease. Lancet 361, 512–519 (2003).

39. Kamphuis, J. B. J., Mercier-Bonin, M., Eutamène, H. & Theodorou, V. Mucus organisation is shaped by colonic content; A new view. Sci. Rep. 7, 1–13 (2017).

40. Azzouz, L. L. & Sharma, S. Physiology, Large Intestine. in *In: StatPearls [Internet]* (StatPearls Publishing, 2025).

41. Chattopadhyay, I. et al. Exploring the Role of Gut Microbiome in Colon Cancer. Appl. Biochem. Biotechnol. 193, 1780–1799 (2021).

42. Chung The, H. & Le, S. N. H. Dynamic of the human gut microbiome under infectious diarrhea. Curr. Opin. Microbiol. 66, 79–85 (2022).

43. Halfvarson, J. et al. Dynamics of the human gut microbiome in inflammatory bowel disease. Nat. Microbiol. 2, 1–7 (2017).

44. Khan, I. et al. Alteration of gut microbiota in inflammatory bowel disease (IBD): Cause or consequence? IBD treatment targeting the gut microbiome. Pathogens 8, 1–28 (2019).

45. Sun, J. Impact of bacterial infection and intestinal microbiome on colorectal cancer development. Chin. Med. J. (Engl). 135, 400–408 (2022).

46. Rath, E. & Haller, D. Intestinal epithelial cell metabolism at the interface of microbial dysbiosis and tissue injury. Mucosal Immunol. 15, 595–604 (2022).

47. Wells, J. M., Rossia, O., Meijerink, M. & Van Baarlen, P. Epithelial crosstalk at the microbiota-mucosal interface. Proc. Natl. Acad. Sci. U. S. A. 108, 4607–4614 (2011).

48. Addie, R. D., Balluff, B., Bovée, J. V. M. G., Morreau, H. & McDonnell, L. A. Current State and Future Challenges of Mass Spectrometry Imaging for Clinical Research. Anal. Chem. 87, 6426–6433 (2015).

49. Sun, Z. et al. Recent strategies for improving MALDI mass spectrometry imaging performance towards low molecular weight compounds. Trends Anal. Chem. 175, 117727 (2024).

50. Guiberson, E. R. et al. Multimodal Imaging of the Murine Gastrointestinal Tract with Retained Luminal Content Supporting Info. J. Am. Soc. Mass Spectrom. 33, 1073–1076 (2022).

51. Liu, X., Flinders, C., Mumenthaler, S. M. & Hummon, A. B. MALDI Mass Spectrometry Imaging for Evaluation of Therapeutics in Colorectal Tumor Organoids. J. Am. Soc. Mass Spectrom. 29, 516–526 (2018).

52. Mirretta Barone, C., et al. Spatially resolved lipidomics shows conditional transfer of lipids produced by Bacteroides thetaiotaomicron into the mouse gut. Cell Host Microbe 32, 1025–1036.e5 (2024).

53. Shi, Q. et al. Investigating the effects of PFOA accumulation and depuration on specific phospholipids in zebrafish through imaging mass spectrometry. Environ. Sci. Process. Impacts 26, 700–709 (2023).

54. Eum, J. H. et al. Long-term liquid nitrogen vapor storage of mouse embryos cryopreserved using vitrification or slow cooling. Fertil. Steril. 91, 1928–1932 (2009).

55. Greene, A. E., Silver, R. K., Krug, M. & Coriell, L. L. Preservation of Cell Cultures by Freezing in Liquid Nitrogen Vapor. Proc. Soc. Exp. Biol. Med. 116, 1–23 (1964).

56. Terracio, L. & Schwabe, K. G. Freezing and Drying of Biological Tissues for Electron Microscopy. J. Histochem. Cytochem. 29, 1021–1028 (1981).

57. Bolon, B. Protocols for Placental Histology. The Guide to Investigation of Mouse Pregnancy (Elsevier, 2014). doi:10.1016/B978-0-12-394445-0.00045-X

58. The Inkscape Team. Inkscape Project. (2003).

59. Bender, K. J. et al. Sample Preparation Method for MALDI Mass Spectrometry Imaging of Fresh-Frozen Spines. Anal. Chem. 95, 17337–17346 (2023).

60. Sumner, L. W. et al. Proposed minimum reporting standards for chemical analysis: Chemical Analysis Working Group (CAWG) Metabolomics Standards Initiative (MSI). Metabolomics 3, 211–221 (2007).

